# Mitochondrial type II NADH dehydrogenase of *Plasmodium falciparum* is dispensable and not the functional target of putative NDH2 quinolone inhibitors

**DOI:** 10.1101/436881

**Authors:** Hangjun Ke, Suresh M. Ganesan, Swati Dass, Joanne M. Morrisey, Sovitj Pou, Aaron Nilsen, Michael K. Riscoe, Michael W. Mather, Akhil B. Vaidya

## Abstract

The battle against malaria has been substantially impeded by the recurrence of drug resistance in *Plasmodium falciparum*, the deadliest human malaria parasite. To counter the problem, novel antimalarial drugs are urgently needed, especially those that target unique pathways of the parasite, since they are less likely to have side effects. The mitochondrial type II NADH dehydrogenase of *P. falciparum*, PfNDH2 (PF3D7_0915000), has been considered a good prospective antimalarial drug target for over a decade, since malaria parasites lack the conventional multi-subunit NADH dehydrogenase, or Complex I, present in the mammalian mitochondrial electron transport chain (mtETC). Instead, Plasmodium parasites contain a single subunit NDH2, which lacks proton pumping activity and is absent in humans. A significant amount of effort has been expended to develop PfNDH2 specific inhibitors, yet the essentiality of PfNDH2 has not been convincingly verified. Herein, we knocked out PfNDH2 in *P. falciparum* via a CRISPR/Cas9 mediated approach. Deletion of PfNDH2 does not alter the parasite’s susceptibility to multiple mtETC inhibitors, including atovaquone and ELQ-300. We also show that the antimalarial activity of the fungal NDH2 inhibitor HDQ and its new derivative CK-2-68 is due to inhibition of the parasite cytochrome *bc*_*1*_ complex rather than PfNDH2. These compounds directly inhibit the ubiquinol-cytochrome *c* reductase activity of the malarial *bc*_*1*_ complex. Our results call into question the validity of PfNDH2 as an antimalarial drug target.

**Importance:** For a long time, PfNDH2 has been considered an attractive antimalarial drug target. However, the conclusion that PfNDH2 is essential was based on preliminary and incomplete data. Here we generate a PfNDH2 KO (knockout) parasite in the blood stages of *Plasmodium falciparum*, showing that the gene is not essential. We also show that previously reported PfNDH2-specific inhibitors kill the parasites primarily via targeting the cytochrome *bc*_*1*_ complex, not PfNDH2. Overall, we provide genetic and biochemical data that help to resolve a long-debated issue in the field regarding the potential of PfNDH2 as an antimalarial drug target.

## Introduction

The mitochondrial electron transport chain (mtETC) is an important, validated drug target in malaria parasites. The mtETC is the primary generator of the electrochemical gradient across the mitochondrial inner membrane. In the asexual blood stages of malaria parasites, however, the only critical function of the mtETC is the continuous reoxidation of ubiquinol to sustain dihydroorotate dehydrogenase (DHODH) activity, which is required for de novo pyrimidine biosynthesis (1). In contrast, in insect stages, mitochondrial oxidative phosphorylation appears to have increased importance (2), likely requiring an intact central carbon metabolism (3) and increased mtETC activity to maintain the electrochemical gradient that drives ATP synthesis. For decades, the mtETC of malaria parasites has attracted major drug development efforts (4), ultimately resulting in antimalarials for clinical use and in preclinical/clinical stages of development. Malarone™, a combination of atovaquone and proguanil, has been used clinically since 2000. Recent drug development efforts focused on the parasite DHODH led to the clinical candidate DSM265, which is currently undergoing Phase II clinical trials (5, 6). ELQ-300, an inhibitor of the Qi site of the *bc*_*1*_ complex (Complex III), has also reached preclinical development (7, 8). This underscores that the essential protein components of the parasite mtETC are attractive antimalarial drug targets.

In the parasite mtETC, there are five dehydrogenases that donate electrons to ubiquinone producing ubiquinol (reduced ubiquinone), which is subsequently oxidized by the *bc*_*1*_ complex (Complex III). These five enzymes include NDH2, malate quinone oxidoreductase (MQO), DHODH, glycerol 3-phosphate dehydrogenase (G3PDH), and succinate dehydrogenase (SDH). As mentioned above, the parasite DHODH is a validated antimalarial drug target. NDH2 has also been considered a promising antimalarial drug target for over a decade (9–12). In general, NADH dehydrogenase is a membrane bound flavoenzyme that catalyzes electron transfer from NADH to quinone producing NAD^+^ and quinol. In human mitochondria, a type I NADH dehydrogenase (Complex I) has 45 subunits and pumps protons across the mitochondrial inner membrane concomitant with electron transfer (13). Mutations of Complex I subunits are responsible for a significant portion of hereditary human respiratory chain disorders (14). In contrast, malaria parasites lack the conventional multi-subunit Complex I. Instead, they have a type II NADH dehydrogenase (NDH2), which is a single subunit, non-proton pumping protein, likely attaching to the mitochondrial inner membrane and facing the mitochondrial matrix. *Toxoplasma gondii*, another apicomplexan parasite, has two isoforms of NDH2, which both face the mitochondrial matrix, catalyzing oxidation of mitochondrial NADH (15). NDH2 is also present in bacteria (16), fungi (17) and plants (18), but not in humans or other mammals.

The absence of NDH2 in humans suggests that the parasite enzyme might be a good antimalarial drug target (9–12). In 1990, Fry and Beesley first measured NADH oxidation activities in isolated mitochondria of malaria parasites (*P. yoelii* and *P. falciparum*) using two spectrophotometric methods (19). Briefly, in the first assay, NADH oxidation was coupled to cytochrome *c* reduction and changes of cytochrome *c* absorption spectrum were measured at a wavelength of 550 nm; in the second assay, NADH oxidation produced NAD^+^, directly leading to a reduced absorption at 340 nm. Using these measurements, Fry and Beesley found that NADH oxidation in the mitochondrial samples was more robust than that of other substrates and was not inhibited by rotenone, a classical Complex I inhibitor. The conclusion was that mitochondria of malaria parasites were able to oxidize NADH, although it was not clear which specific enzyme(s) were responsible or which pathway(s) were involved. In 2006, Biagini *et al.* also observed significant NADH oxidation activity (direct assay at 340 nm) in *P. falciparum* extracts (9). Biagini *et al.* used atovaquone and potassium cyanide to block the activities of Complexes III and IV individually, leading them to conclude that the observed NADH oxidation was due to PfNDH2 (9). However, with the use of total cell extracts containing various NADH dependent enzymes, it seems questionable to attribute all the observed NADH oxidation activity to PfNDH2 alone (9, 12). Coincidentally at that time, the ubiquinone analogue HDQ (1-hydroxy-2-dodecyl-4(1H) quinolone) was found to be a potent inhibitor of the fungal NDH2 in *Yarrowia lipolytica* (20). Later HDQ was shown to be highly effective against *P. falciparum* and *T. gondii* parasites (10). Based on these results (9, 10, 12), it became widely accepted that PfNDH2 could be an attractive antimalarial drug target. As a result, a significant drug discovery campaign based on high throughput screening was undertaken to seek HDQ-like inhibitors to specifically inhibit PfNDH2 (21–23), yielding the lead compound, CK-2-68 (22). Recently, the crystal structure of PfNDH2 was resolved via x-ray crystallization (24), which could further encourage drug development efforts towards PfNDH2 using approaches based on in silico docking and structure activity relationships of PfNDH2 inhibitors.

The rationale for targeting PfNDH2 for antimalarial drug development has, however, been controversial (25, 26). The fact that the entire mtETC in asexual blood stages could be functionally bypassed by expression of the heterologous yDHODH from *Saccharomyces cerevisiae* to support pyrimidine biosynthesis in the presence of mtETC inhibition raised the likelihood that PfDHODH is the only essential enzyme among the five mitochondrial dehydrogenases that donate electrons to ubiquinone (1). The yDHODH transgenic parasites can be grown continuously under a high atovaquone pressure (100 nM) (1, 27). Under such conditions, the *bc*_*1*_ complex is fully inhibited, which prevents the reoxidation of ubiquinol by the mtETC and, therefore, should block the turnover of all subsequent quinone-dependent dehydrogenases, implying that PfNDH2, as well as PfG3PDH, PfMQO, and PfSDH, are not required for growth. Interestingly, drug development research towards PfNDH2 inhibitors did not appear to slow down (21–23) after these results were reported (25), nor even when the type II NADH dehydrogenase in the rodent malaria parasite *P. berghei*, PbNDH2, was genetically ablated in 2011 (28). Very recently, selection of resistant *P. falciparum* parasites by treatment with CK-2-68 and RYL-552, reported “PfNDH2 specific” inhibitors, generated mutations in the mtDNA encoded *cyt b* locus, while no mutations were found in PfNDH2 (29); these data strongly suggests that CK-2-68 and RYL-552 exert their antimalarial activity by inhibiting the parasite *bc*_*1*_ complex, not PfNDH2, in contrast to previous suggestions (21–23). However, in the absence of specific genetic data on the essentiality of PfNDH2, the importance of PfNDH2 in *P. falciparum* has not been settled definitively.

Here, we successfully knocked out PfNDH2 in *P. falciparum* using a CRISPR/Cas9 based approach, which should put to rest the question of the validity of PfNDH2 as an antimalarial drug target. We found that deletion of PfNDH2 did not alter the parasite’s susceptibility to major mtETC inhibitors and, further, that HDQ and CK-2-68 kill malaria parasites by directly inhibiting the parasite cytochrome *bc*_*1*_ complex.

## Materials and Methods

### 1. Parasite maintenance and transfection

*P. falciparum* D10 is the wildtype (WT) parasite line used in this study. D10attB-yDHODH was generated previously (27), which expresses the yeast DHODH gene of *Saccharomyces cerevisiae*. Parasites were cultured with RPMI 1640 medium (Invitrogen by Thermo Fisher Scientific) supplemented with 5g/L Albumax I (Invitrogen), 10 mg/L hypoxanthine, 2.1 g/L sodium bicarbonate, HEPES (15 mM), and gentamycin (50 μg/ml). Cultures were maintained in human red blood cells (Type O, Interstate Blood Bank, Tennessee) and kept in a CO_2_/O_2_ incubator filled with a low oxygen mixture (5% O_2_, 5% CO_2_, and 90% N_2_). Ring stage parasites with 5% parasitemia were electroporated with plasmid DNA in cytomix buffer using a Bio-Rad gene pulser. Drug medium was added 48 h post electroporation. For hdhfr (human dihydrofolate dehydrogenase) selectable marker, 5 nM WR99210 was used.

### 2. Plasmid construction

1. Removal of yDHODH from the pAIO pre-gRNA construct. The pre-gRNA construct pAIO was generously provided by Dr. Josh Beck (30); the plasmid contains yDHODH and *Streptococcus pyogenes* Cas9 coding sequences (CDS) connected by a 2A “self-cleaving” peptide. To remove yDHODH, pAIO was digested with BamHI and BglII to release the entire sequence of yDHODH and the first 250 bp of Cas9, since there is no unique restriction site between the two genes that could be used to release the yDHODH CDS alone. The first 250 bp of Cas9 were amplified from the original pAIO vector with primers P1 and P2, which include short homologous sequences that match the ends of the pAIO vector after its digestion with BamHI and BglII. The PCR product and the digested vector were then joined together using NEBuilder® HiFi DNA Assembly (New England Biolabs®, Inc). A colony PCR was performed to screen colonies using primers P1 and P2. Positive clones were grown up, and their plasmid DNAs were digested with BamHI and BglII to confirm the loss of yDHODH. The positive plasmids were then sequenced using a primer upstream of Cas9 (P3) to confirm the intactness of Cas9. These procedures yielded the pre_gRNA construct without yDHODH, namely pAIO-yDHODH(-).
2. PfNDH2 KO construct. PfNDH2 (PF3D7_0915000) is 1602 bp long with no introns. We cloned the 5’ and 3’ homologous regions of PfNDH2 into a pCC1 vector bearing the hdhfr selectable marker (31). The 5’HR (934 bp) was amplified with primers P4 and P5 and cloned into pCC1 digested by NcoI and EcoRI. Subsequently, the 3’HR (936 bp) was amplified with primers P6 and P7 and cloned into the vector digested by SpeI and SacII. After cloning, both 5’HR and 3’HR were sequenced (Genewiz LLC). The KO construct was named 5’3’PfNDH2_pCC1. Maxi prep DNA of 5’3’PfNDH2_pCC1 (Qiagen) was digested with HincII overnight to linearize the vector before transfections.
3. Guide RNA constructs. The sequence between the 5’HR and 3’HR of PfNDH2 (490 bp) was submitted to the gRNA design tool (http://grna.ctegd.uga.edu/) to seek potential gRNAs. From the list of candidates, three sequences were chosen based on their high scores and zero off-target predictions. For each of these sequences, a pair of complementary oligonucleotides (60 or 61 bp) was synthesized and annealed in a mixture of NEB Buffers 2 and 4 by heating to 95°C for 5 minutes, then slowly cooling to room temperature. The vector, pAIO-yDHODH(-), was digested with BtgZI and joined with the annealed oligonucleotide pair by gene assembly (New England Biolabs®, Inc), yielding a pAIO-yDHODH(-)-gRNA construct. Other gRNA cloning procedures followed our published protocol (32).

Primers used for cloning procedures are listed below.

P1 (Remove yDHODH-F), 5’-ATACCTAATAGAAATATATCAGGATCCAAAAATGGACAAGAAGTACAGCATCG;
P2 (Remove yDHODH-R), 5’-CCATCTCGTTGCTGAAGATC;
P3 (Remove yDHODH-chk), 5’-GTATATTTTAAACTAGAAAAGGAATAAC;
P4 (KO-5fF), 5’-GACCATGGATATCAAAAAATAATGCAGTAAAATGC;
P5 (KO-5fR), 5’-CCGAATTCTGAACCTAGGATTATAATCTTTTCTTTTC;
P6 (KO-3fF), 5’-CTACTAGTGTCGAAGTTACCGCAGAATTTG;
P7 (KO-3fR), 5’-AACCGCGGTCTTAATAAAATCGATGAAAAAATGGAACC;
P8 (gRNA1-F), 5’-CATATTAAGTATATAATATTgAATGTACCACTACATAAACAGTTTTAGAGCTAGAAATAGC;
P9 (gRNA1-R), 5’-GCTATTTCTAGCTCTAAAACTGTTTATGTAGTGGTACATTcAATATTATATACTTAATATG;
P10 (gRNA2-F), 5’-CATATTAAGTATATAATATTgCATGTAGCTGTTGTAGGAGGGTTTTAGAGCTAGAAATAGC;
P11 (gRNA2-R), 5’-GCTATTTCTAGCTCTAAAACCCTCCTACAACAGCTACATGcAATATTATATACTTAATATG;
P12 (gRNA3-F), 5’-CATATTAAGTATATAATATTgTTATTTAATTATAGCTGTAGGTTTTAGAGCTAGAAATAGC;
P13 (gRNA3-R), 5’-GCTATTTCTAGCTCTAAAACCTACAGCTATAATTAAATAAcAATATTATATACTTAATATG;
P14 (gRNA1-N20), 5’-AATGTACCACTACATAAACA;
P15 (gRNA2-N20), 5’-CATGTAGCTGTTGTAGGAGG;
P16 (gRNA3-N20), 5’-TTATTTAATTATAGCTGTAG;
P17 (N20CheckR), 5’-ATATGAATTACAAATATTGCATAAAGA;
P18 (5fchk), 5’-GAACTATACATCTATAAAGCATTAC;
P19 (3fchk), 5’-GAAAAAAGAAGCACATATATATATAT;
P20 (hDHFR-F), 5’-ATGCATGGTTCGCTAAACTGCATC;
P21 (hDHFR-R), 5’-ATCATTCTTCTCATATACTTCAAATTTGTAC.

### 3. Assessing parasite growth

PfNDH2 KO and D10 WT lines were synchronized several times by alanine (0.5 M, pH 7.6 with 10 mM HEPES) treatment. On day 0, parasites were inoculated into a 24 well plate with each well containing 2.5 ml of culture at 1% parasitemia and 3% hematocrit. Cultures were fed daily and split every two days. At each split (1:5), a sample of the parasitized RBCs was pelleted and fixed with 4% paraformaldehyde at 4°C overnight. After all samples were collected and fixed, they were washed with 1× PBS and stained with SYBR green I at 1:1000 (Catalog S7567, Life technologies by ThermoFisher Scientific). The samples were washed with PBS three times and analyzed on a C6 Flow Cytometer (BD). A total of 250,000 events were collected for each sample. Unstained infected RBCs and stained uninfected RBCs served as negative controls for gating. Growth curves were drawn using Graphpad Prism 6.

### 4. Growth inhibition assays using ^3^H-hypoxanthine incorporation

Inhibitor compounds were diluted by a series of three-fold dilutions in 96 well plates in low hypoxanthine medium (2.5 mg/L). Parasites were washed three times with low hypoxanthine medium, supplemented with fresh blood sufficient to make 1% parasitemia and re-suspended in the proper volume of low hypoxanthine medium to make a suspension with 3% hematocrit. Aliquots of the diluted culture were added to the 96-well plates containing the inhibitor dilution series. After 24 h incubation, 10 μl of 0.5 μCi ^3^H-hypoxanthine was added to each well and the plates were incubated for another 24 h. After a total of 48 h incubation, the parasites were lysed by freezing-and-thawing, and nucleic acids were collected onto a filter using a cell harvester (Perkin Elmer). Radioactivity was counted using a Topcount scintillation counter (Perkin Elmer). Data were analyzed and graphed using Graphpad Prism 6.

### 5. Ubiquinol-cytochrome *c* reduction assay

Mitochondria of ΔPfNDH2 and D10 WT were individually isolated using a method published previously (32, 33). Briefly, a large volume of parasite culture of each line (~2 liter) was lysed with saponin (0.05%) and disrupted in a N_2_ cavitation chamber (Parr 4639 Cell Disruption Bomb) in an isotonic mitochondrial medium. The total parasite lysate was spun down at 900 *g* for 6 min to remove large debris, and the cloudy supernatant was passed through a MACS CS column (Miltenyi Biotec) in a Vario MACS magnetic separation apparatus to remove most of the hemozoin. The eluted light yellow material was pelleted at 23,000 × *g* for 20 min at 4 °C, and the pellet was re-suspended in buffer and stored at −80 °C. The cytochrome *c* reductase activity of the *bc*_1_ complex was measured with a modification of previous methods (32–34). The assay volume was 300 μl, containing mitochondrial proteins (~5-10 μl mitochondrial preparation), 100 μM decylubiquinol, 75 μM horse heart cytochrome *c* (Sigma-Aldrich), 0.1 mg/ml n-docecyl-β-D-maltoside, 60 mM HEPES (pH 7.4), 10 mM sodium malonate, 1 mM EDTA, and 2 mM KCN, and was incubated at 35°C in a stirred cuvette. Reduction of horse heart cytochrome *c* was recorded at 550 nm with a CLARITY VF integrating spectrophotometer (OLIS, Bogart, GA). A Bio-Rad colorimetric assay was used to measure protein concentrations of all mitochondrial samples.

### 6. NADH-cytochrome *c* reductase assay

Assay conditions were similar to those described above for the ubiquinol-cytochrome *c* reduction assay. The 300 μL assay mix contained 5-10 μl of mitochondrial proteins, 50 μM horse heart cytochrome *c*, 60 mM HEPES (pH 7.4), 10 mM sodium malonate, 1 mM EDTA, 2 mM KCN and 300 μM NADH. The assay buffer contained no detergent, since it was reported that detergents heavily interfere with assays of NADH oxidation (35).

## Results

### PfNDH2 is not essential in asexual blood stages of *Plasmodium falciparum*

Transcriptomics data indicate that the type II NADH dehydrogenase in *P. falciparum* (PF3D7_0915000) is expressed in the asexual blood stages (PlasmoDB.org). It has been shown that the leader sequence of PfNDH2 was able to target GFP into the mitochondrion (36), suggesting that PfNDH2 is a mitochondrial enzyme. To further confirm that, we genetically tagged PfNDH2 with 3× HA and the tagged PfNDH2 was localized to the parasite mitochondrion by immunofluorescence assays (37). To assess the essentiality of PfNDH2, we employed the CRISPR/Cas9 DNA repair technique. A KO plasmid vector was constructed (Figure 1A, Materials and Methods), containing a 5’HR (homologous region) mostly upstream of the gene’s coding sequence (CDS) (outside) and a 3’HR near the end of the CDS (inside). The 3’HR was chosen from the coding region to circumvent inclusion of overly high AT content in the KO vector. Three gRNA sequences targeting the PfNDH2 gene were individually cloned into a modified pre-gRNA-Cas9 plasmid construct, from which yDHODH had been removed (Materials and Methods). Previous studies have shown that expression of yDHODH in malaria parasites renders the entire mtETC nonessential by providing a metabolic bypass for pyrimidine biosynthesis (1). Therefore, to assess the essentiality of PfNDH2 in the context of a normal mtETC, we removed yDHODH from the gRNA vectors. The KO plasmid was linearized by restriction digestion and transfected into D10 parasites together with the three circular gRNA vectors (Materials and Methods). Viable transgenic parasites were observed under WR99210 selection three weeks post transfection. As shown in Figure 1B, a diagnostic PCR revealed that PfNDH2 was disrupted. We then tightly synchronized both ΔPfNDH2 and WT lines and examined the growth rates over 4 intraerythrocytic developmental cycles (IDCs) via flow cytometry. As shown in Figure 1C, ΔPfNDH2 and WT parasites grew equally well over this time period. The ΔPfNDH2 KO line was further maintained in culture for over one month, and no growth defects were noticeable (data not shown). Deletion of PfNDH2 also did not appear to affect parasite health and morphology (Figure 1D). Collectively, our data indicate that PfNDH2 is not essential in asexual blood stages of *P. falciparum*, consistent with the KO study carried out previously in the rodent malaria parasite, *P. berghei* (28). These results argue against the long-held assumption that PfNDH2 is an attractive drug target (9–12).

**Figure 1.**
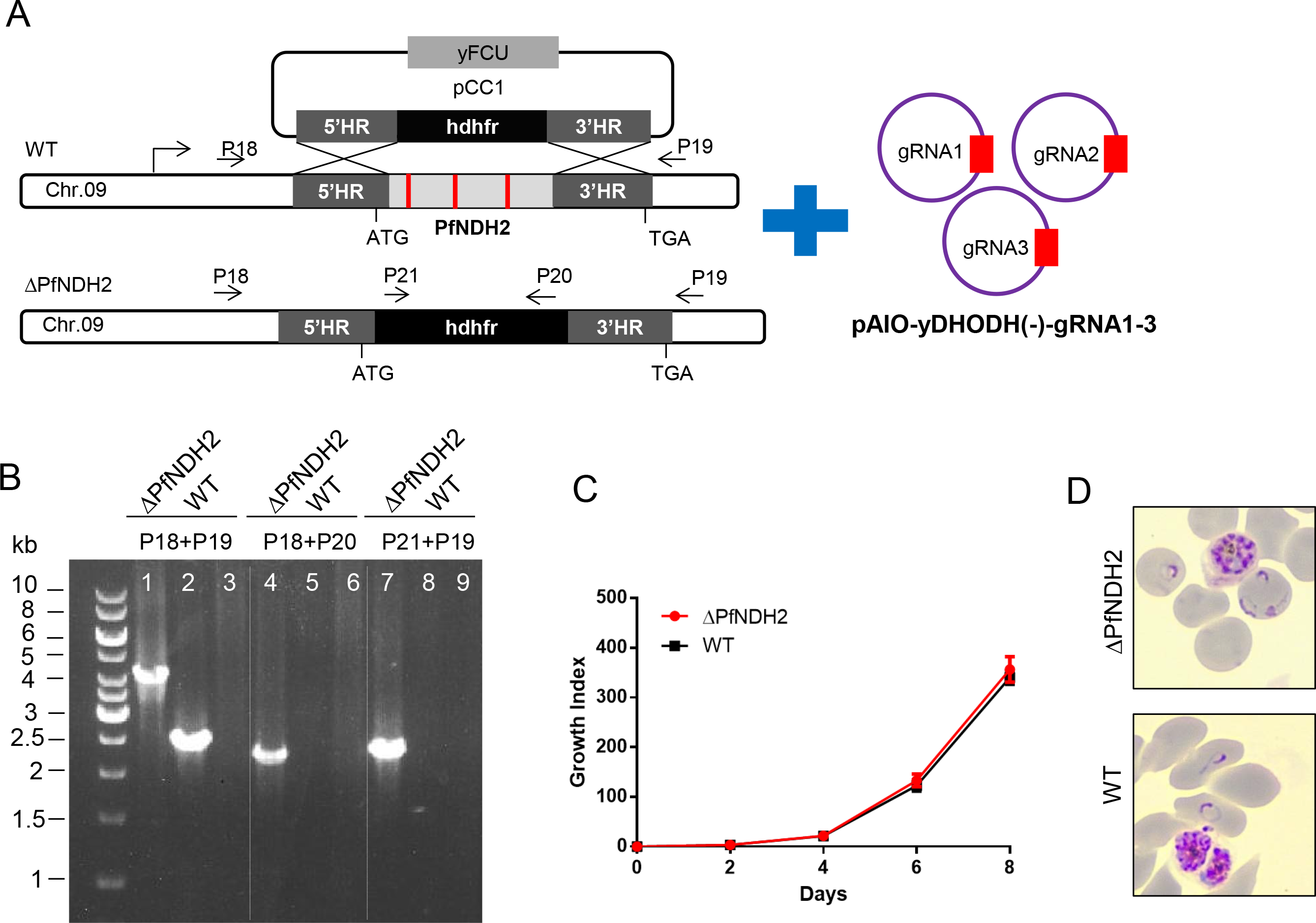
Disruption of the type II NADH dehydrogenase of *P. falciparum* does not affect growth in asexual blood stages. (A) A schematic diagram depicts the genetic deletion of a large segment of PfNDH2 via CRISPR/Cas9-assisted homologous recombination. (B) A diagnostic PCR confirming the genotype of the ΔPfNDH2 parasite. Primer positions are shown in (A). In ΔPfNDH2, a 4.1 kb knockout band (Lane 1), a 2.2 kb 5’ integration band (Lane 4) and a 2.5 kb 3’ integration band (Lane 7) were detected. In WT, only a 2.5 kb band was detected (Lane 2) whereas no 5’ integration (Lane 5) or 3’ integration (Lane 8) was observed. Lanes 3, 6, and 9 were negative controls with no DNA in PCR reactions. (C) A growth curve of the ΔPfNDH2 parasite determined by SYBR green staining and flow cytometry analysis (Materials and Methods). (D) The ΔPfNDH2 parasite is morphologically healthy. Giemsa stained thin smears of WT and ΔPfNDH2 cultures are shown displaying schizont and ring forms.

### The ΔPfNDH2 parasite is equally susceptible to mtETC inhibitors

The healthy growth of the ΔPfNDH2 parasites *in vitro* (Figure 1) suggests that the parasite mtETC remains functionally competent in the absence of PfNDH2. To challenge the KO parasites, we exposed them to mtETC inhibitors in growth inhibition assays, measured as ^3^H-hypoxanthine incorporation. As shown in Figure 2, in comparison to the WT, the ΔPfNDH2 parasites were equally sensitive to atovaquone (a Q_o_ site inhibitor of the *bc*_*1*_ complex) and ELQ-300 (a Q_i_ site inhibitor) (8). Thus, these data suggest that deletion of PfNDH2 has little effect on the sensitivity of asexual parasites to downstream inhibitors of the mtETC. The loss of NDH2, thus, does not appear to affect the function of the remainder of the mtETC. As noted previously, HDQ and its newer derivative CK-2-68 were considered to be PfNDH2 specific inhibitors (21–23) or, more recently, dual-targeting inhibitors of cytochrome *bc*_*1*_ as well as PfNDH2 (38). In that case, HDQ and CK-2-68 would be expected to lose potency in the ΔPfNDH2 parasite, since the putative primary target is absent. However, HDQ and CK-2-68 were still highly potent in the KO parasite (Figure 2), suggesting that HDQ and CK-2-68 primarily target another site than PfNDH2.

**Figure 2.**
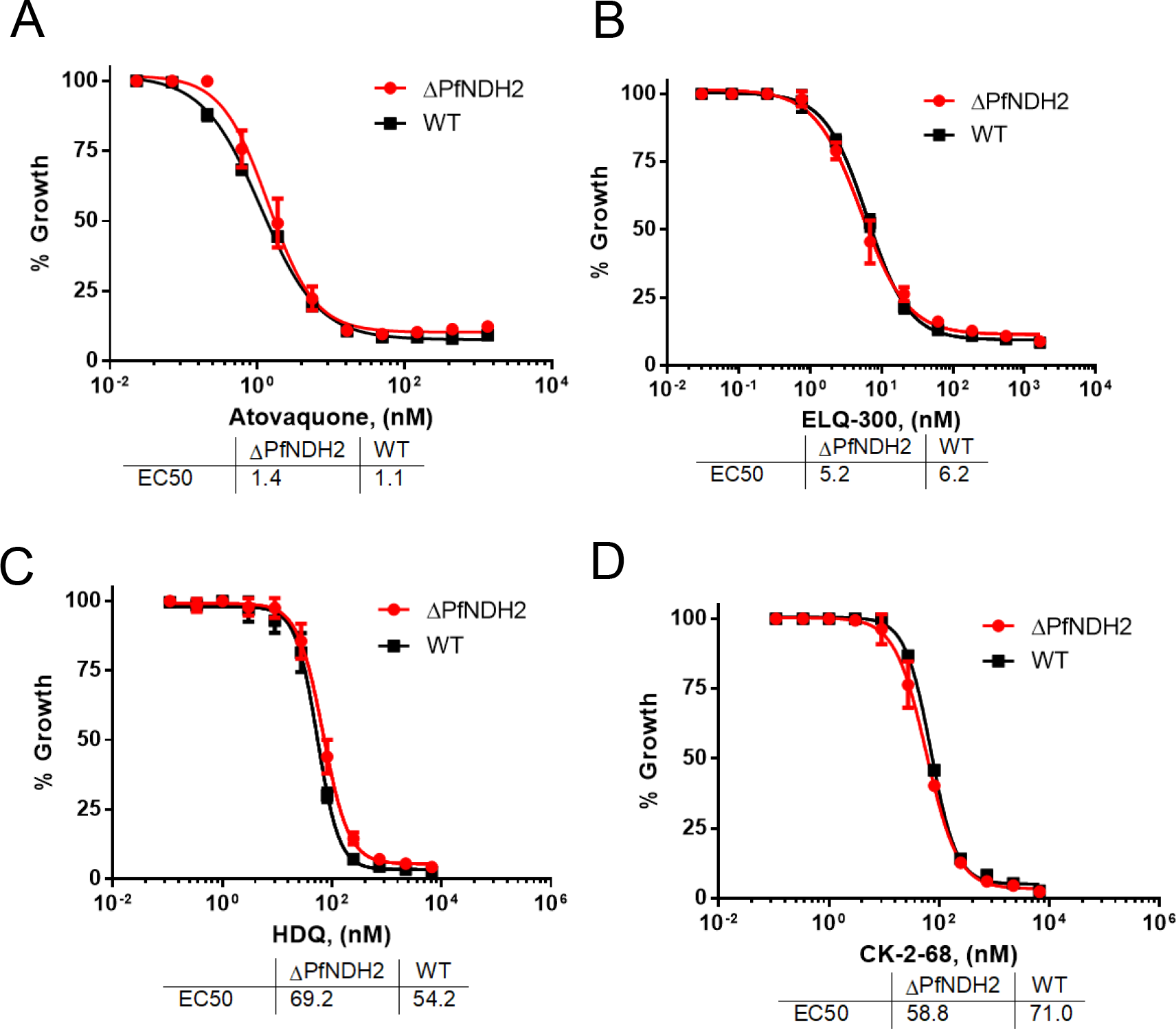
Deletion of PfNDH2 does not affect sensitivity to mitochondrial electron transport chain inhibitors. ^3^H-hypoxanthine incorporation assays were performed in the ΔPfNDH2 and WT parasites challenged with atovaquone (A), ELQ-300 (B), HDQ (C) and CK-2-68 (D). Data shown is a representative of n≥3 replicates.

### The *bc*_*1*_ complex of the mtETC is the target of HDQ and CK-2-68

Our data above suggests that HDQ and CK-2-68 target an activity other than PfNDH2 (Figure 2). Since HDQ and CK-2-68 are ubiquinone analogs, we suggest that they kill malaria parasites by targeting the *bc*_*1*_ complex, although Vallieres *et al.* and Biagini *et al.* previously suggested that HDQ and CK-2-68 had a dual effect on both PfNDH2 and the *bc*_*1*_ complex (38, 39). To distinguish between these alternatives, we performed growth inhibition assays in the yDHODH transgenic parasite line using HDQ and CK-2-68 in combination with proguanil. As shown previously, expression of the yDHODH gene bypasses the need for mtETC function in asexual parasites (1). The yDHODH transgenic parasites have become a handy tool to examine whether a compound targets the mtETC, as all mtETC inhibitors suffer a large loss of potency in the yDHODH background, which applies to both *bc*_*1*_ inhibitors and PfDHODH inhibitors (40). Further, a low concentration of proguanil (1 μM) can restore sensitivity to *bc*_*1*_ inhibitors in yDHODH transgenic parasites (1), but not for PfDHODH inhibitors. As a control, we showed that yDHODH parasites were fully resistant to atovaquone but became fully sensitive in the presence of 1 μM proguanil (Figure 3). Upon inhibition by atovaquone, the yDHODH parasites lose their primary source of ΔΨ_m_ generation, conveyed by the *bc*_*1*_ complex and cytochrome *c* oxidase of the mtETC, and become hypersensitive to proguanil, which targets a secondary generator of ΔΨ_m_ (1). Using this system, we tested the HDQ and CK-2-68 sensitivity of the yDHODH parasites with and without 1 μM proguanil. As shown in Figure 3, yDHODH parasites were highly resistant to HDQ and CK-2-68, as expected; upon proguanil treatment, the yDHODH parasites regained sensitivity to these compounds. HDQ and CK-2-68, thus, behaved in a very similar manner to atovaquone against the yDHODH transgenic parasites, indicating that HDQ and CK-2-68 target the *bc*_*1*_ complex. These results are consistent with a previous report that found that parasites grow normally in the presence of 10 μM HDQ when expressing the yDHODH gene (41). Furthermore, recent chemical mutagenesis experiments using CK-2-68 generated mutations that were all in the *cyt b* locus, rather than in PfNDH2 (29). Collectively, these results indicate that HDQ and CK-2-68 are potent cytochrome *bc*_*1*_ inhibitors.

**Figure 3.**
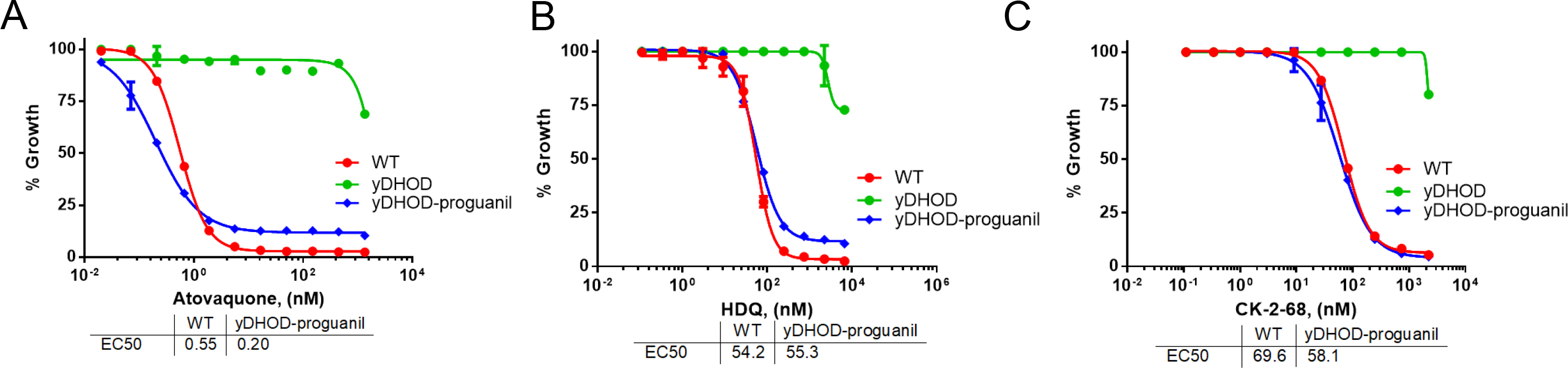
HDQ and CK-2-68 target the *bc*_*1*_ complex. D10attB-yDHODH transgenic parasites were challenged with atovaquone (A), HDQ (B) and CK-2-68 (C) with and without proguanil (1 μM). Growth was measured using ^3^H-hypoxanthine incorporation assays. With an EC_50_ of ~60 μM, 1 μM of proguanil had no effect on WT parasites (1).

### HDQ and CK-2-68 directly inhibit the enzymatic activity of the *bc*_*1*_ complex

In addition to growth inhibition assays as described above (Figures 2 and 3), we also directly investigated the effect of HDQ and CK-2-68 on the enzymatic activity of the *bc*_*1*_ complex in a preparation enriched in parasite mitochondria using a spectrophotometric assay (Materials and Methods) (33). As shown in Figure 4, HDQ and CK-2-68 inhibited the ubiquinol-cytochrome *c* reductase activity in the mitochondria of ΔPfNDH2 and WT in a dose dependent manner. Importantly, the inhibitory potency of HDQ and CK-2-68 were equally robust in two types of mitochondria from WT and ΔPfNDH2, respectively. This provides further evidence that the antimalarial mode of action of HDQ and CK-2-68 arises from inhibition of the *bc*_*1*_ complex, rather than PfNDH2.

**Figure 4.**
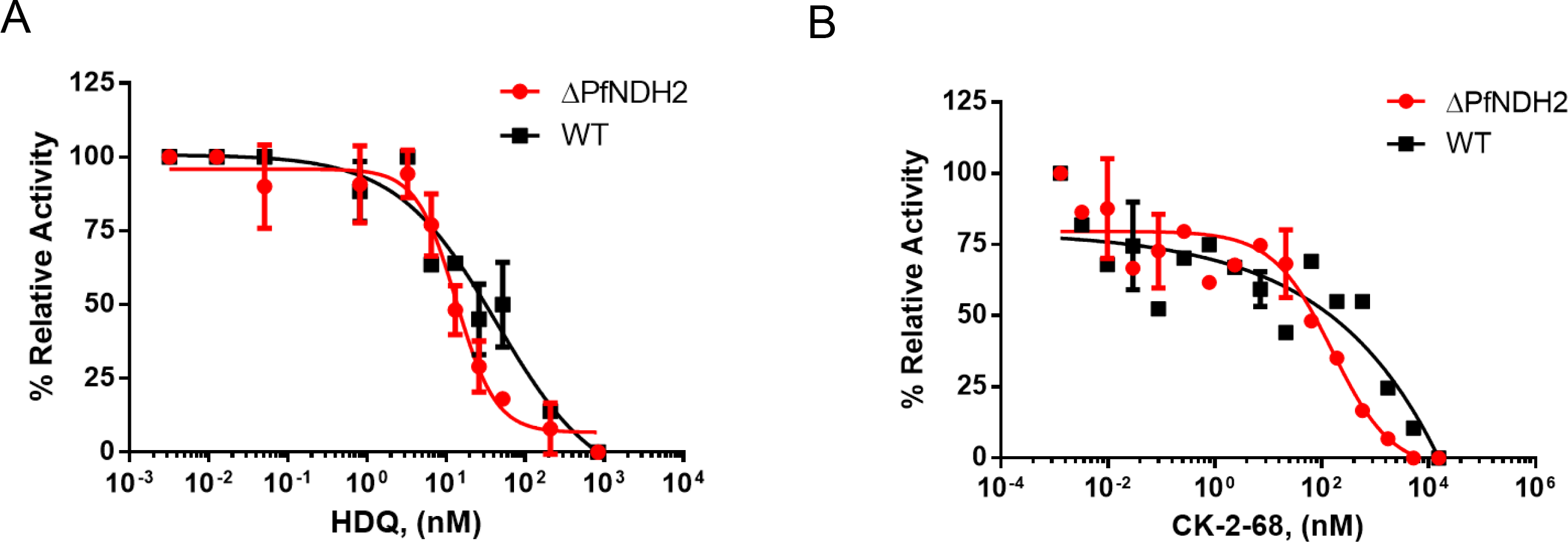
HDQ and CK-2-68 directly inhibit mitochondrial ubiquinol-cytochrome *c* reductase activity *in vitro*. In each measurement, the same amount of mitochondria of KO or WT (10 μl of sample) was used. Reduction of cytochrome *c* was followed spectrophotometrically at 550 nm (Materials and Methods). The rates of reduction in the presence of various concentrations of inhibitors were normalized to that of no drug controls (average of 4-5 replicates), resulting in relative activity (%). (A) Inhibition by HDQ. Data shown is plotted from n=3 biological replicates. (B) Inhibition by CK-2-8. Data shown is plotted from n=2 biological replicates.

### In vitro measured NADH linked cytochrome *c* reductase activity is likely non-biological

Previously data from Fry and Beesley (19) revealed a relatively strong NADH-cytochrome *c* reductase activity in parasite mitochondrial preparations. Interestingly, the activity was not inhibited by rotenone (80 μM) or antimycin A (a Qi inhibitor at 20 μM) (19). Rotenone insensitivity suggested that malaria parasites lack a conventional multi-subunit Complex I, which was later interpreted as evidence that the type II NADH dehydrogenase was essential (9). While antimycin A did not inhibit the NADH-cytochrome *c* reductase activity in Fry and Beesley’s mitochondrial preparations, it did inhibit the cytochrome *c* reductase activity when other mitochondrial substrates were used, such as α-glycerophosphate and succinate (19). In addition, antimycin A kills malaria parasites in whole cell assays with an EC_50_ of 13 nM (42). Thus, the provenance of the apparent NADH-cytochrome *c* reductase activity observed in mitochondrial preparations has been an unsettled issue. In intact parasites, NADH oxidized by NDH2 is presumed to pass electrons to ubiquinone, which are then transferred on to the *bc*_*1*_ complex, cytochrome *c*, cytochrome *c* oxidase and, finally, to O_2_. If the *in vitro* assay were replicating the initial steps of the *in vivo* pathway, we should observe a much diminished NADH-cytochrome *c* reductase activity in the ΔPfNDH2 parasites since PfNDH2, missing in the knockout parasite, is the only known enzyme donating electrons from NADH to the mtETC in the parasites (15). As shown in Figure 5, however, deletion of PfNDH2 had no effect on NADH-cytochrome *c* reductase activity, suggesting that this *in vitro* assay is likely non-physiological. Further, in both ΔPfNDH2 and WT mitochondria, NADH-cytochrome *c* reductase activity was not inhibited by a mix of malaria parasite specific *bc*_*1*_ inhibitors, including atovaquone (62 nM), ELQ-300 (62 nM), and HDQ (3,100 nM), each at equal or greater than 100× EC_50_ (Figure 5). Thus, our data are consistent with the earlier observation that antimycin A failed to inhibit the NADH-cytochrome *c* reductase assay (19) and suggest that the *in vitro* NADH-cytochrome *c* reductase activity is likely non-enzymatic (see Discussion).

**Figure 5.**
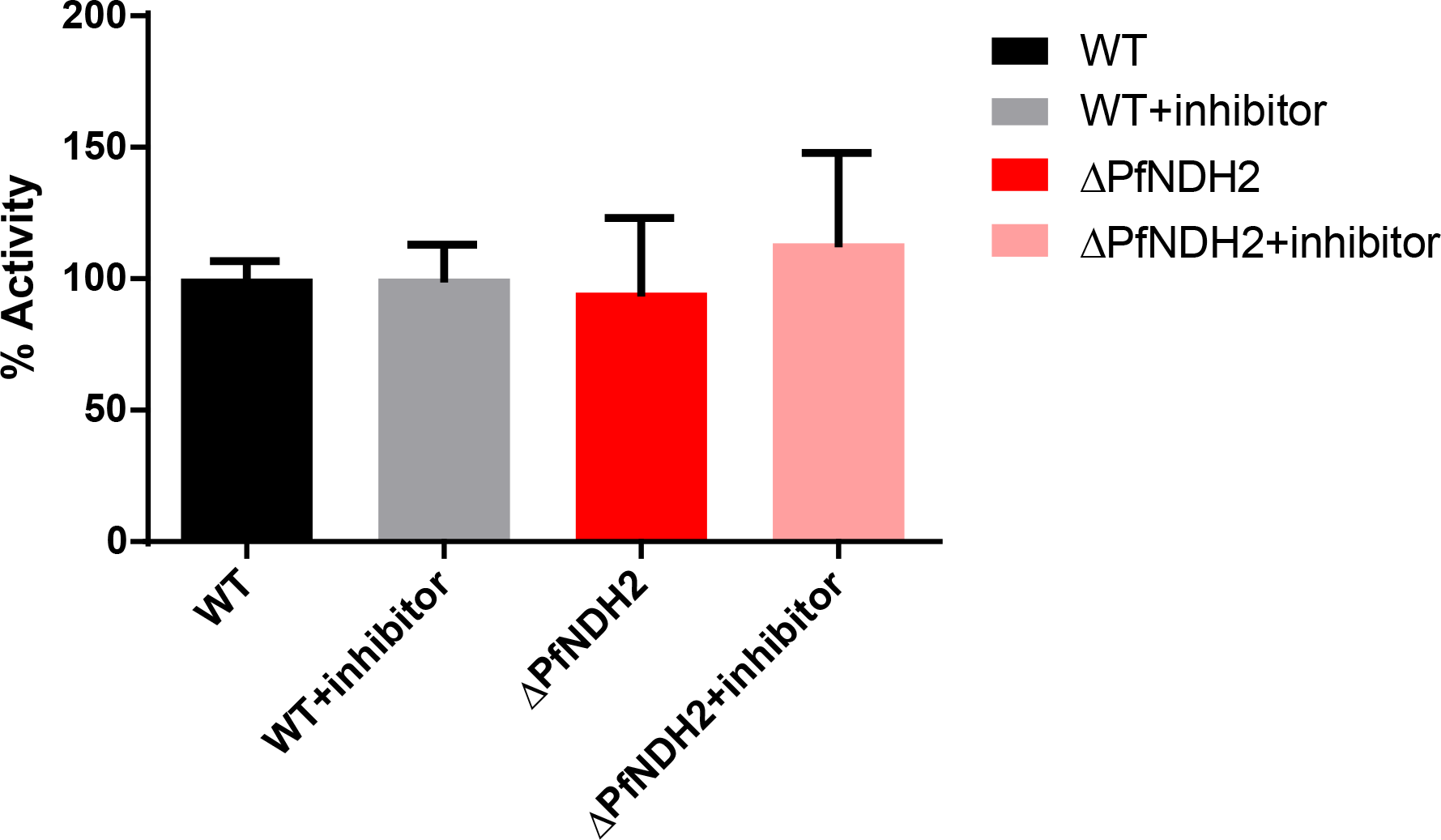
NADH-linked cytochrome *c* reductase activity in the *in vitro* assay is not dependent on PfNDH2 or mtETC. In each measurement, 10 μl of mitochondrial sample was used. Rates of cytochrome reduction were measured without and with addition of *bc*_*1*_ inhibitors (atovaquone (62 nM), ELQ-300 (62 nM) and HDQ (3,100 nM)). Data shown is mean ± s.d. of n=3 replicates.

## Discussion

A common strategy for developing antimicrobial drugs is to target divergent proteins of the microbe to circumvent potential toxicity against the host. Proteins unique to microbes are even more interesting as their inhibitors would potentially have little to no side effects in the host. The type II NADH dehydrogenase is present in malaria parasites but not in humans; thus, it has been considered an attractive prospective drug target for a long time (9, 11, 43). However, a unique protein may not necessarily be an essential one. A valid drug target should normally be essential to the pathogen in order that its inhibition will arrest growth and/or kill the pathogen. Initial failures to disrupt the NDH2 gene in *P. falciparum* parasites suggested that the gene might be essential (35, 37). On the other hand, data on the effect of mtETC inhibitors in yDHODH transgenic parasites (1) and the reported knockout of NDH2 in *P. berghei* (28) suggested that PfNDH2 should be dispensable in asexual parasites (as described above). Without conclusive data, however, it remains a long-debated issue in the field whether PfNDH2 is a good antimalarial drug target. In this report, we have provided strong evidence indicating that PfNDH2 is dispensable in asexual blood stages and, therefore, is unlikely to be an effective antimalarial drug target. We note that our knockout result is consistent with the recent genetic screen of *P. falciparum* growth phenotypes, in which a PiggyBac transposon insertion was recovered in the CDS of PfNDH2, suggesting non-essentiality of the gene (44).

Our results suggest that the parasite mtETC is functionally intact in the absence of PfNDH2. Not only is the growth of the KO line closely similar to that of the WT parental line (Figure 1C), but the response to cytochrome *bc*_*1*_ inhibitors is virtually identical (Figure 2). Evidently, in the ΔPfNDH2 parasites, the other ubiquinone-dependent dehydrogenases-MQO, SDH, G3PDH, and DHODH—supply sufficient ubiquinol to maintain adequate function of the mtETC during asexual development. DHODH is essential for the parasite’s pyrimidine de novo synthesis pathway, since malaria parasites cannot salvage pyrimidine precursors; the other dehydrogenases, however, may be functionally redundant as electron donors to the mtETC. We have previously carried out a comprehensive genetic and biochemical study in the TCA cycle of *P. falciparum* (3). In the asexual blood stages, the main carbon source of the TCA is glutamine, rather than glucose. KDH (alpha-ketoglutarate dehydrogenase) is the entry point of glutamine derived carbons into the TCA cycle. KDH converts alpha-ketoglutarate to succinyl-CoA, with the concomitant reduction of NAD^+^ to NADH, and it is likely that KDH is the principal producer of NADH in the mitochondrial matrix due to a relatively large TCA flux observed with labeled glutamine (3). Yet, neither KDH nor the TCA flux contributed by glutamine is essential to the parasite in asexual blood stages, which is consistent with the non-essential nature of PfNDH2 as a consumer of NADH.

Although HDQ and CK-2-68, and probably other related derivatives (29) do not primarily target PfNDH2 in parasites, as shown by our results, they are potent antimalarial compounds via inhibition of the parasite *bc*_*1*_ complex. Importantly, HDQ and CK-2-68 retained their potency in atovaquone resistant parasites (38, 39). Experiments with yeast *cyt b* mutants suggested that HDQ likely bound to the Qi site of *bc*_*1*_ complex, whereas atovaquone is a Qo site inhibitor (38). CK-2-68, on the other hand, is likely to be a Qo site inhibitor, but, nevertheless, exhibited no cross resistance with atovaquone (29). Biagini *et al.* have developed additional quinolone derivatives with more favorable pharmacological properties that were predicted to bind at the Qo site (39). Combinations of non-cross resistant *bc*_*1*_ inhibitors may be effective at slowing the development and spread of resistance, since strong resistance mutations in *cyt b* may exert a significant survival fitness cost (45), including blocking transmission (46). Thus, the development of additional antimalarial candidates targeting the *bc*_*1*_ complex may facilitate the future development of effective combination therapies. Indeed, atovaquone and ELQ-300, Q_o_ and Q_i_ inhibitors respectively, were recently shown to be a highly effective and synergistic antimalarial combination (47).

The results of our attempts to measure *in vitro* NADH-cytochrome *c* reductase activity spectrophotometrically provide a cautionary tale for the design and interpretation of assays involving the oxidation of NADH, a reactive reductant. Neither elimination of PfNDH2 nor strong inhibition of the cytochrome *c* reductase activity of *bc*_*1*_ affected the observed reaction (Fig. 5), implying that the reaction does not proceed through the mtETC. Fry and Beasley apparently observed the same phenomenon when they measured apparent NADH-cytochrome *c* reductase activity in *Plasmodium* mitochondria with and without antimycin A, a general Qi site inhibitor of the *bc*_*1*_ complex (19). Given the report that detergents (which form micelles) accelerate NADH oxidation (35), we speculate that it may be the presence of mitochondrial phospholipid membranes in the mitochondrial samples that produce this effect. Cytochrome *c* is known to bind to phospholipids head groups (48), so mitochondrial particles could provide a surface that concentrates cytochrome *c* for reaction with NADH (a trimolecular reaction, requiring 2 cytochromes *c* to oxidize one NADH, as it is a 2-electron reductant). The non-enzymatic reaction may also be facilitated by the relatively high concentration of cytochrome *c* used in spectrophotometric assays (50-100 μM). At any rate, our results demonstrate that the apparent robust NADH-cytochrome *c* activity that has been reported in *Plasmodium* mitochondrial preparations *in vitro* is not an indication of high NADH dehydrogenase activity in intact parasites.

## Acknowledgements

We thank Dr. Kristin D. Lane and Dr. Thomas E. Wellems of NIH/NIAID for kindly providing the CK-2-68 compound. The Cas9-gRNA construct was generously provided by Dr. Josh R. Beck (now at Iowa State University) and Dr. Daniel E. Goldberg from Washington University St. Louis. This research was supported by an NIH grant (ABV, R01 AI028398) and an NIH K22 award (HK, K22 AI127702). MKR receives grants from NIH (R01, AI100569), Veterans Affairs Merit Review Program Award (i01 BX003312) and DOD (Log# PR130649; Contract # W81XWH-14-1-0447).

## Author contribution statements

HK and ABV designed the outline of the project. HK produced the PfNDH2 KO parasite and characterized its genotypes and phenotypes. SMG made the KO construct. SD and MWM measured NADH cytochrome *c* reductase activities *in vitro*. JMM performed growth inhibition assays. SP, AN and MKR synthesized HDQ and ELQ-300. HK wrote the manuscript which was modified by all other authors.

